# Does larval ability to modulate body buoyancy explain successful colonization of freshwater environments by diadromous gobies?

**DOI:** 10.1101/2023.07.27.550194

**Authors:** Yumeki Oto, Katsutoshi Watanabe

**Author notes:** Corresponding author: Y Oto.

## Abstract

Salinity is an environmental factor that strongly characterizes the habitat use patterns of aquatic organisms. However, knowledge is biased toward the effect of differences in osmotic pressure among salinity habitats; how ambient specific gravity (SG) differences determine species distribution is scarcely understood. Diadromous fish, which migrate between marine and freshwater habitats, may encounter this SG problem when they are unexpectedly landlocked in or colonize freshwater areas with low environmental SG. This is particularly serious for planktonic larval fish, which must maintain neutral buoyancy for foraging and passive locomotion, although their swimbladders are generally underdeveloped. Then, we hypothesized that the SG problem limits the establishment of freshwater resident populations in marine-originated diadromous fishes. To test this hypothesis, the SG modulation ability of newly hatched larvae was compared among three closely related diadromous goby species in *Gymnogobius*, one of which has freshwater resident populations. The aquarium experimental results did not support that only the species deriving freshwater residents can maintain neutral buoyancy even in freshwater conditions; that is, all three species could modulate their body SG almost equally to those of both fresh and sea waters. This suggests that the ability to maintain neutral buoyancy in freshwater had been pre-adaptively acquired prior to larval freshwater colonization. On the other hand, it is highly noteworthy that the early larvae of the target group maintained neutral buoyancy in various SG environments using swimbladders, which is the first such evidence in teleosts.

## 1. Introduction

Salinity is an environmental factor that critically affects the survival of aquatic organisms and strongly determines their distribution patterns (Schultz and McCormick, 2013). It has been well documented that failure of osmotic regulation quickly kills teleost fish, often preventing them from entering their non-natal salinity environments (Parry, 1966; Schultz and McCormick, 2013). Another important physical factor that varies with ambient salinity is specific gravity (SG), which can also affect fish survival (Holliday, 1969). Particularly in planktonic larvae, the failure to maintain neutral buoyancy results in higher energy consumption to maintain their position in the water column (Czesny et al., 2005; Woolley and Qin, 2010) and morphological abnormalities, such as lordotic deformities (Chatain, 1989; Schwebel et al., 2018). An ecological study revealed that the failure of swimbladder inflation negatively affects foraging and predator avoidance in the larval yellow perch *Perca flavescens*, reducing growth and survival rates (Czesny et al., 2005). Thus, modulation of larval body SG is necessary for growth and survival. However, information is quite limited about how strongly the differences in SG among ambient waters determine the pattern of salinity habitat use by larval fish.

Diadromous fish, which regularly migrate between sea and freshwater areas throughout their lives, are prone to situations requiring them to acclimate to different environmental SG. Particularly, this SG problem could be serious if larval fish encounter a change in ambient salinity because the regulation range of their body SG is generally narrow owing to the undeveloped swimbladder (Battaglene and Talbot, 1990; Kitajima et al., 1993; Woolley and Qin, 2010; He et al., 2022). Several studies have reported that marine planktonic larvae of diadromous species cannot maintain neutral buoyancy well under freshwater conditions with a low SG (approximately 1.000). For example, the monk goby *Sicyopterus japonicus* exhibits 1.034–1.036 in body SG during the marine larval period and fails to maintain neutral buoyancy under freshwater conditions (Iida et al., 2010, 2013). Similarly, the marine larvae of several other goby species exhibit higher body SG (1.017–1.041) than SG of ambient low-salinity brackish water (approximately 1.007), suggesting a poor acclimation ability to low SG conditions (Iida et al., 2017). Therefore, for acclimatization to low-salinity habitats, planktonic larvae that usually inhabit the sea would have to innovate their physiological systems to decrease their body SG.

Some diadromous species establish freshwater resident (FWR) populations in lakes and streams (Delgado and Ruzzante, 2020). This is an ecologically and evolutionarily significant phenomenon that triggers the further adaptive evolution of diadromous species to local environments, sometimes resulting in the derivation of strictly freshwater species (Lee and Bell, 1999; Waters et al., 2020). For marine-originated diadromous species, larval survival under freshwater conditions is the key to the establishment of FWR populations. Among the several types of diadromy, amphidromy is the most speciose and provides the clearest example in which larval physiology limits the establishment of FWR populations because most amphidromous species enter the sea only in the larval stage, when physiological flexibility is generally low (McDowall, 2007; Augspurger et al., 2017; Delgado and Ruzzante, 2020). Among amphidromous fishes, the FWR form has been observed only in a limited number of species or groups (Tsunagawa and Arai, 2008; Augspurger et al., 2017; Oto et al., 2022). However, no study has yet determined the larval fish characteristics that are critical for the occurrence of FWR populations in amphidromous species.

Groups in which the presence or absence of the FWR type differs among closely related species are useful for identifying key adaptive traits for freshwater colonization by minimizing noise due to non-targeted physiological and ecological differences. The genus *Gymnogobius* (Gobiidae), a marine-originated goby group, includes diadromous species both with and without FWR populations, in addition to marine, brackish water, and strictly freshwater species (Stevenson, 2002). A clade in *Gymnogobius* includes three amphidromous (*G. petschiliensis*, *G. opperiens*, and *G. urotaenia*) and one strictly freshwater species (*G. isaza*) (Aizawa et al., 1994; Harada et al., 2002; Ito et al., 2022). Among the three amphidromous species, only *G. urotaenia* contains several FWR populations in lakes and streams, whereas the others spend their larval periods in the sea (Umino et al., 2015; Oto et al., 2022). A previous study demonstrated the relative superiority of the osmoregulatory performance of *G. urotaenia* larvae under freshwater conditions (Oto et al., 2022). However, this may not be sufficient to fully explain the life-history variations among *Gymnogobius* species because of some osmoregulatory flexibility also in the other two species. The ability to regulate body SG in larval fish is another important trait in their survival in freshwater environments. Therefore, the difference in environmental SG between freshwater and seawater habitats is expected to be a strong limiting factor in establishing FWR populations (SG hypothesis). In our hypothesis, *G. urotaenia* larvae were predicted to adjust the SG of their body almost equal to that of freshwater conditions, even just after hatching, whereas *G. petschiliensis* and *G. opperiens* larvae could not adjust their body SG well to freshwater and hence exhibited higher body SG than freshwater environmental SG.

To test our hypothesis, larval body SG was experimentally compared among three *Gymnogobius* species reared under freshwater and seawater conditions. Furthermore, the relationship between body SG and swimbladder size was examined to confirm whether the swimbladder plays a crucial role in SG regulation. Finally, the swimming layers of the three species were recorded under each salinity condition to confirm the effect of body SG regulation on buoyancy. The results of the present study are expected to provide insights into the potential role of the SG modulation ability, an under-highlighted physiological trait, in the evolution of life history through novel niche exploitation.

## 2. Materials and methods

### 2.1 Target species and sampling

This study targeted three amphidromous *Gymnogobius* species, *G. petschiliensis*, *G. opperiens*, and *G. urotaenia*, which are distributed across the Japanese Archipelago, Korean Peninsula, and Yellow Sea coast (Stevenson, 2002; Ishino, 2005; Ito et al., 2022). All three species usually spawn in freshwater stream areas, migrate downstream to the sea immediately after birth, drift along the coast for 2–3 months during the larval stage, and return to streams when juveniles, forming large schools (Dotu, 1955; Harada, 2002). However, only *G. urotaenia* has established freshwater resident (FWR) populations in lakes and streams in several regions (Umino et al., 2015; Oto et al., 2022). No FWR populations have been reported for *G. petschiliensis* or *G. opperiens*.

To acquire newly hatched larvae, egg clutches of the target species were sampled from two or three populations per species inhabiting five rivers and one lake located on the Pacific and Sea of Japan coasts in Japan (population ID PI and PA: *G. petschiliensis*; OO and ON: *G. opperiens*; UE, UA, and UB: *G. urotaenia*; 3–9 clutches each; Fig. 1). These samples were common to those used in our previous study (Oto et al., 2022). The UA and UB of *G. urotaenia* were determined to be the FWR populations by otolith microchemical analysis of the parental males of the egg clutches used in this study (Oto et al., 2022). The water temperature and salinity at each site were recorded using an electronic salinometer (ES-71; Horiba, Ltd., Kyoto, Japan). The water temperature at the coastal sea surface was investigated using data from the Japan Meteorological Agency (https://www.data.jma.go.jp/gmd/kaiyou/data/db/kaikyo/daily/sst_HQ.html) to determine the laboratory temperature.

**Fig. 1.**
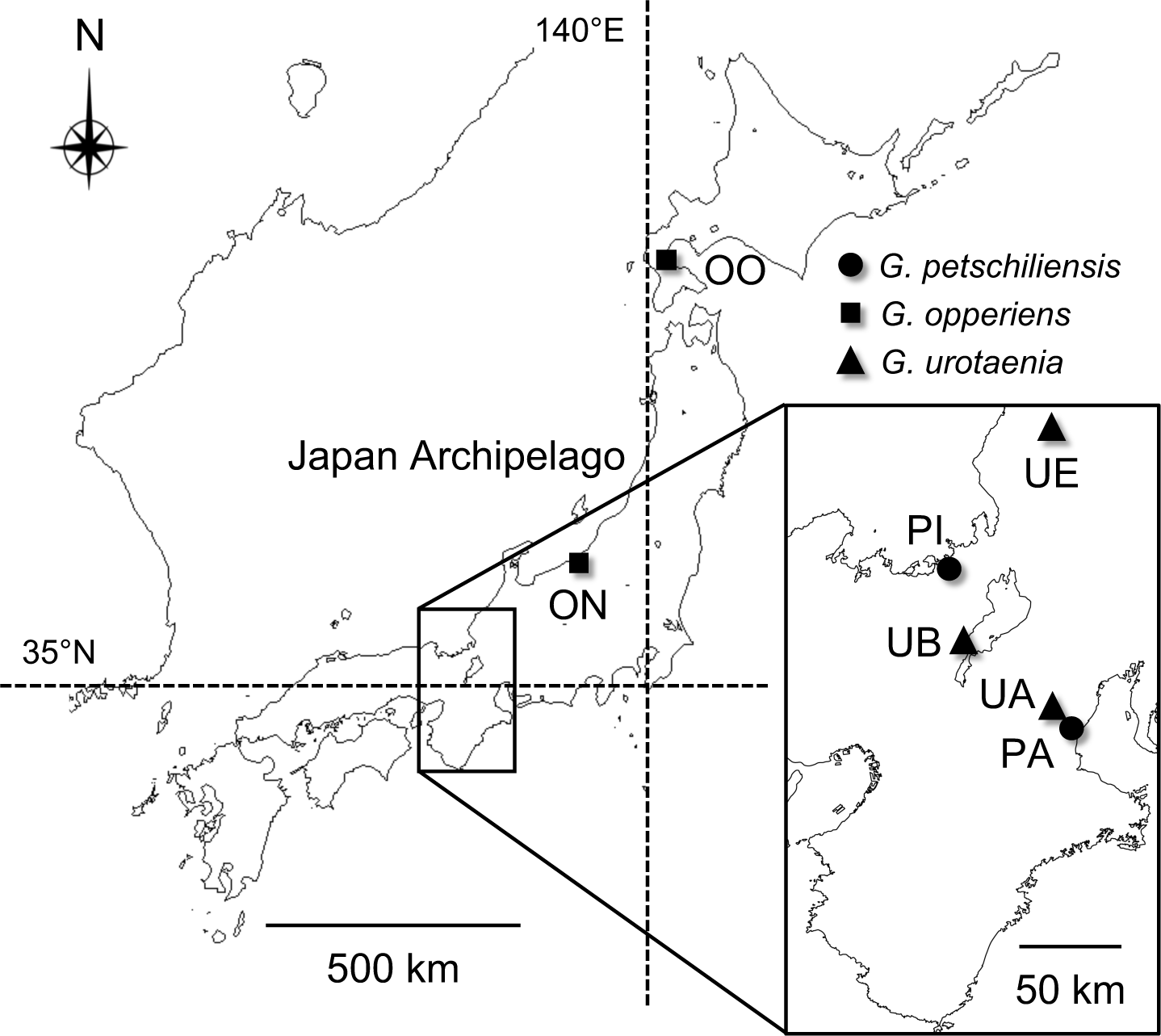
Location of the sampling points of the *Gymnogobius* gobies. *G. petschiliensis*: PI, Inokuma River; PA, Ano R.; *G. opperiens*: OO, Okutsunai R.; ON, Nigorisumi R.; *G. urotaenia*: UE, Eiheiji R.; UA, Ano R.; UB, Lake Biwa.

### 2.2 Larval rearing

We conducted a rearing experiment on newly hatched larvae from 8–14 clutches of each species to compare their ability to regulate the body specific gravity (SG) among the three *Gymnogobius* species. Egg clutches attached to stones (spawning beds in the wild) were transported in buckets filled with aerated river waters to the laboratory located at Kyoto University (Sakyo, Kyoto, Japan), where the temperature was maintained at 13– 18 °C (approximately intermediate between those of sampling sites and the sea surface). When more than half of the eggs hatched in each clutch 0–9 days after sampling, exactly 18 individuals of larval fish were randomly selected and carefully transferred into 2 L experimental tanks filled with aerated freshwater (salinity 0.1; freshwater-reared group) and seawater (salinity 34.5–35.5; seawater-reared group) using 10 ml pipettes. Salinity was measured using an electronic salinometer (ES-71). This transition timing was determined based on the estimated time for the newly hatched larvae to reach the sea in the wild; all clutches (except the landlocked UB population) were found at sites 0.1–30 km from the river mouths. The rearing water comprised tap water dechlorinated with sodium thiosulfate, followed by addition of an artificial seawater medium (SEALIFE; Nihonkaisui Co. Ltd., Tokyo, Japan) to make the seawater. The larvae were fed 0.1 g of frozen rotifers *Branchionus plicatilis* (Kyorin Co., Ltd., Himeji, Japan) per tank twice daily. Live rotifers were not supplied because their survivability and mobility were negatively affected by freshwater conditions (e.g., Imentai et al., 2019).

### 2.3 Measurement of body specific gravity

To measure the body SG, two larvae were randomly sampled from each rearing bottle 3 and 7 days after hatching, except when insufficient individuals survived. The protocol for measuring SG was based on Tsukamoto et al. (2009) and Iida et al. (2010). The body SG was measured at daytime under the same laboratory conditions as for larval rearing. Using the artificial seawater medium and sodium polytungstate (Sometu, Berlin, Germany), a series of solutions with SG values ranging from 1.0010 to 1.0490 was prepared at intervals of 0.0015 (32 steps). Solutions with SG values ranging from 1.0010 (freshwater) to 1.0250 (seawater) were prepared by adding a precisely weighed artificial seawater medium to the ion exchange water. Solutions with >1.0250 in SG were prepared by adding sodium polytungstate to avoid larval death due to excessive hyperosmotic stress, which was also a concern in Tsukamoto et al. (2009). Two sets of 32 test solutions were prepared. To prevent inaccurate measurements due to active swimming, 0.03% 2-phenoxyethanol (Wako Pure Chemical Industries, Ltd., Osaka, Japan) was dissolved in the measurement solutions as an anesthetic.

The measurement protocol of larval body SG consisted of two steps. First, the larval body was prewashed using one of the two sets of solutions (prewashing solutions) to minimize the contamination of rearing freshwater or seawater. In this prewashing step, the larval SG was roughly estimated by transferring them to several prewashing solutions. Subsequently, the main measurement was conducted using the other set of solutions (main solutions). In this experimental step, the larvae were observed for approximately 30 seconds to determine whether their body SG was in a neutral buoyancy state in a solution with the same SG as that used in the previous step. If the neutral buoyancy of the anesthetized larvae was maintained, the SG of the solution was adopted as that of the larval body specimen. When neutral buoyancy was not perfect, the SG of the test solution that most effectively reduced the floating or sinking was regarded as the larval body SG. All larvae were transferred between tanks or bottles with a few milliliters of rearing water after being carefully absorbed using a 10 mL pipette. The main test solutions used for the measurements were replaced with those stored in a refrigerator at 4 °C before starting similar measurements on later days.

### 2.4 Measurement of swimbladder

The larvae were photographed under anesthesia after the SG measurement to quantify the size of the larval swimbladder. After placing the fish with their body sides on top, the horizontal cross-sectional area of the swimbladder and notochord length were measured using the measurement function of a digital microscope (VW-9000; Keyence Corporation, Osaka, Japan). The data were not used if an anesthetized individual died because of fragility during filming (judged by observing a beating heart).

### 2.5 Observation of swimming layer

Larval swimming layers in freshwater and seawater were observed under a laboratory condition in a tank measuring 12 × 9 × 15 cm (length × width × depth). The back and bottom of the tank were opaque blue, and the front was transparent. A horizontal line was drawn on the front side of the tank to enable the observer to distinguish the upper- and lower-layer sections of the water column (that is, 7.5 cm in depth for each layer section). All surviving 4-day-old larvae (1–16 individuals per rearing bottle) were carefully transferred to the experimental tank filled with rearing water, placed in a compartment covered with a dark curtain, and illuminated with aquarium lighting (1070 lm × 2). After the larvae were acclimated in the experimental tank for 30 minutes, an observer counted the number of individuals swimming in the upper-layer section and those that were floating (not sinking to the bottom) in the tank every 30 seconds for a total of 21 times during a 10-minute period. After the experiment, all larvae were returned to the rearing bottles and observed to swim normally. This 10-minute observation experiment was performed once for each rearing bottle.

### 2.6 Statistical analysis

The analyses were performed based on the following four comparison groups: *G. petschiliensis*, *G. opperiens*, stream populations of *G. urotaenia*, and lake-locked *G. urotaenia* (UB). These four comparison groups of the three species are hereafter referred to as four “species” for convenience. The analyses of body SG and swimbladders were not conducted on the 7-day-old *G. opperiens* because the number of surviving individuals was insufficient (*n* = 3 for each salinity). All except for the lake-locked *G. urotaenia* included two stream populations. The R software (version 4.1.3; R Core Team 2022) was used for all analyses.

First, an intraspecific comparison of the body SG was conducted each for the 3- and 7-day-old larvae between the freshwater- and seawater-reared groups using the Mann–Whitney U test. In this analysis, data from two populations of each species (except UB) were pooled. The calculated *P*-values were corrected using the Holm method (Holm, 1979) according to the number of intraspecific comparison pairs (four and three pairs for 3- and 7-day-old larvae, respectively).

Next, the size of the larval swimbladders was compared between the freshwater- and seawater-reared groups for each of the four species. As swimbladders were also considered to be correlated with body size, a multiple regression analysis was conducted by running a linear mixed model (LMM). In this model, the response variable was the cross-sectional area of swimbladders. The predictor variables were salinity (categorical; freshwater or seawater) and notochord length (mm^2^). The IDs of the larval population and clutch were also included as random effects. The LMM was generated and run using the lmer function in R (the lme4 package: Bates et al. 2007). The significance of the coefficient was tested using the lmerTest package (Kuznetsova et al., 2017), and the *P*-values were corrected using the Holm method (Holm, 1979) to account for the repetition of intraspecific comparisons.

Finally, the larval swimming layers were compared between salinity conditions and among species. For each rearing bottle with 1–16 fish, the numbers of individuals swimming in the upper-layer section and floating in the tank 5 and 10 minutes after the start of the experiments were selected as representative values for this analysis. As the variable for their swimming layers, individuals swimming at the upper-layer section or floating were assigned as “1”, whereas the remaining, swimming at the lower layer or sinking to the bottom, were assigned as “0”. Next, to test whether salinity conditions predicted the binary variables for the two swimming indices for each species, generalized linear mixed models (GLMMs) were generated for each observation time (5- and 10-minute time points) using the lmer function (Bates et al. 2007). In the models, the IDs of the population and clutch were included as random effects. The response 0/1 variable for the individual swimming layers was assumed to follow a binomial distribution. The GLMMs were run independently for each of the four species and the two indices of the swimming layer. Next, data from the four species were combined into a single dataset to test interspecific differences in the swimming layers. GLMMs similar to those used for intraspecific comparisons were run using this dataset after adding species as a predictor variable (four-level categorical). The significance of the coefficients of salinity condition and species was tested using the lmerTest package (Kuznetsova et al., 2017). Because the intraspecific comparison tests were repeated four times, and the interspecific comparisons were conducted for six pairs of the four species for each observation time, the *P*-values were corrected according to the test repetition using the Holm method (Holm, 1979).

## 3. Results

### 3.1 Body specific gravity

In the larvae of all target *Gymnogobius* species, body specific gravity (SG) was significantly lower in the freshwater condition than in the seawater at both 3 and 7 days of age (Mann–Whitney U test, Holm-corrected *P* < 0.007; Fig. 2). Moreover, the body SG of all species and populations was almost equal to that of freshwater and seawater when reared in the freshwater and seawater conditions, respectively (Means in freshwater: 1.001–1.011; seawater: 1.025–1.033; Fig. 2). Note that the body SG of some freshwater-reared larvae may have been smaller than 1.001 because the larval body sometimes slowly rose to the surface of the water column with an SG of 1.001. The minimum recorded SG value was 1.001, which could have caused a slight underestimation of the SG modulation ability in freshwater environments.

**Fig. 2.**
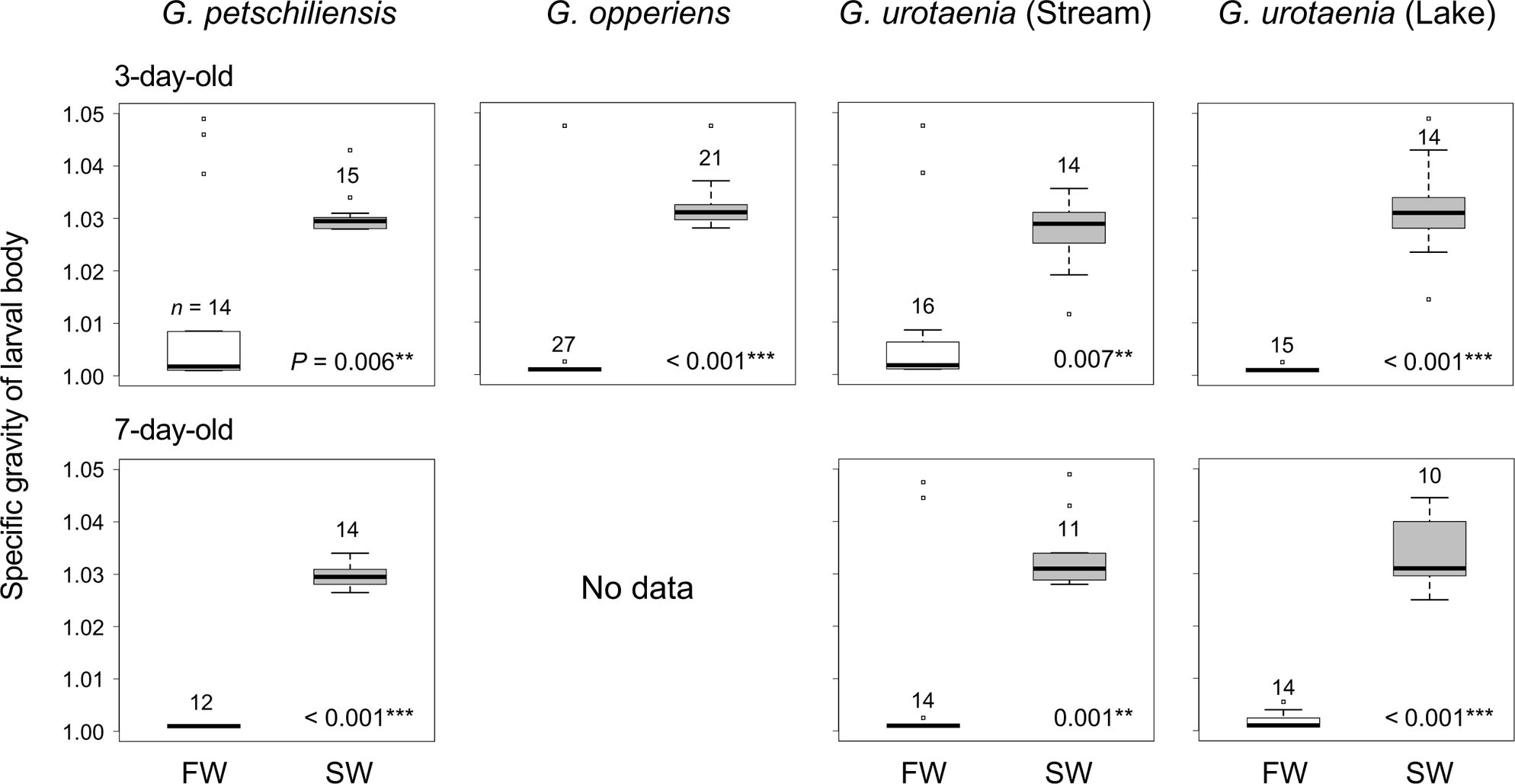
Comparison of the larval body specific gravity (SG) of *Gymnogobius* larvae between freshwater (FW) and seawater (SW) conditions or among the three species/populations. The box plots colored in grey and white show the interquartile range of larval SG under FW and SW conditions, respectively. Open plots represent the outliers, which fall more than 1.5 times the interquartile range above the third quartile or below the first quartile. The *P*-values in the lower right corner of frames indicate the probability that the SG was not different between FW and SW conditions in each species at the same age (\**P* < 0.05, \*\**P* < 0.01, \*\*\**P* < 0.001). The sample size for each measurement is shown above the box plots.

### 3.2 Swimbladder size

The swimbladders of most larval individuals had already inflated 3 days after hatching in all species (Fig. 3A, B). The cross-sectional areas of the larval swimbladders were significantly larger in the freshwater condition than in the seawater, regardless of species and age (LMM, freshwater coefficient = 0.036–0.060, Holm-corrected *P* < 0.018; Fig. 3A, Table 1). The residual error structures of swimbladder size (mm^2^) did not significantly deviate from the Gaussian distribution in each salinity group for each species (Shapiro–Wilk test, *W* = 0.915–0.959, *P* = 0.092–0.588, Fig. S1), justifying the adoption of the linear mixed model.

**Fig. 3.**
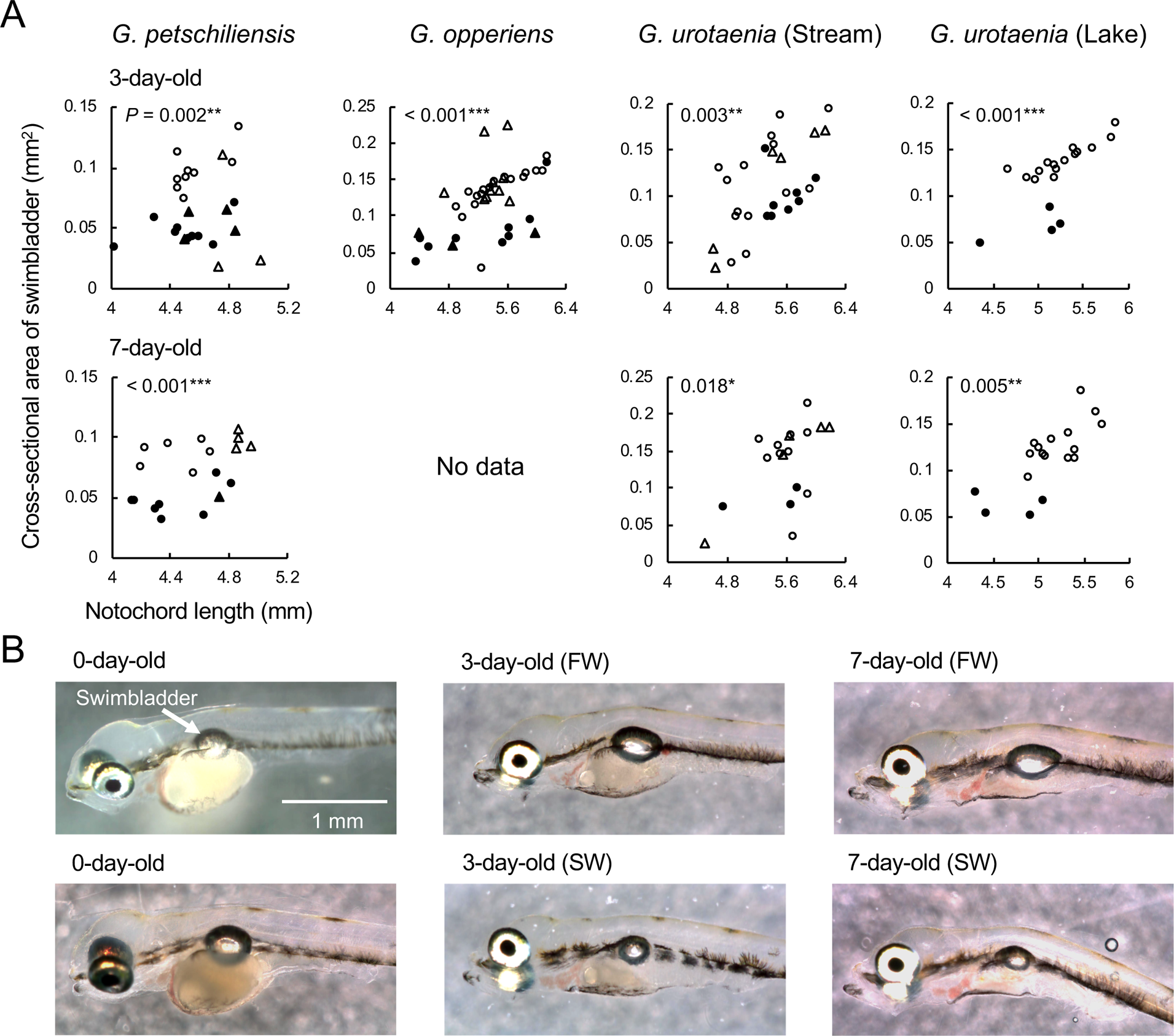
Changes in swimbladder size between freshwater (FW) and seawater (SW) conditions in larval fish of the *Gymnogobius* species. Panel A indicates the relationship between swimbladder and body sizes in FW and SW conditions. The plot color and shape indicate the salinity group (white: FW, black: SW) and population (circle: PA, OO, UA; triangle: PI, ON, UE). The *P*-values in the upper left corner of frames indicate the probability that the swimbladder size was not different between FW and SW conditions in each species at the same age (\**P* < 0.05, \*\**P* < 0.01, \*\*\**P* < 0.001). Panel B shows selected photographs of swimbladders of the 0-, 3-, and 7-day-old larvae from the lake *G. urotaenia* (UB) population.

**Table 1.** Results of the linear mixed models (LMMs) for assessing the difference in swimbladder size between freshwater (FW) and seawater (SW) groups of the three *Gymnogobius* species.

| Species | Age | Coefficient (FW–SW) | Standard error | Degree of freedom | *t* | *P* | Holm corrected *P* |
| --- | --- | --- | --- | --- | --- | --- | --- |
| *G. petschiliensis* | 3-day-old | 0.036 | 0.010 | 21.5 | 3.472 | 0.002** | 0.002** |
|  | 7-day-old | 0.039 | 0.005 | 14.8 | 7.988 | < 0.001*** | < 0.001*** |
| *G. opperiens* | 3-day-old | 0.053 | 0.011 | 35.5 | 4.943 | < 0.001*** | < 0.001*** |
| *G. urotaenia* (River) | 3-day-old | 0.045 | 0.012 | 18.4 | 3.786 | 0.001** | 0.003** |
|  | 7-day-old | 0.060 | 0.022 | 11.8 | 2.756 | 0.018* | 0.018* |
| *G. urotaenia* (Lake) | 3-day-old | 0.060 | 0.005 | 16.0 | 11.026 | < 0.001*** | < 0.001*** |
|  | 7-day-old | 0.053 | 0.013 | 8.5 | 4.218 | 0.003** | 0.005** |

**Note 1:** The formula of LMM was as follows: Cross-sectional area of swimbladder (mm<sup>2</sup>) ~ Salinity group + Square of notochord length (mm<sup>2</sup>). Salinity group was a categorical variable (i.e., FW or SW). Population and parent IDs of each larva were also included as random effects.
**Note 2:** \**P* < 0.05, \*\**P* < 0.01, \*\*\**P* < 0.001.

### 3.3 Swimming layer

For both the ratios of individuals swimming in the upper layer and those floating in the tank, no significant time-dependent changes were found in any species (binomial GLMM for each species, Holm-corrected *P* > 0.229; see Figs. S2, S3). The ratio of individuals swimming in the upper-layer section significantly varied only in stream *G. urotaenia* with salinity at both 5 and 10 minutes after the start of the observation (11.7% in freshwater vs. 29.9% in seawater; binomial GLMM, seawater coefficient = 1.032– 1.327, Holm corrected *P* < 0.017; Fig. 4). Similar significant salinity-dependent variation was found in *G. opperiens* and lake *G. urotaenia* at the 10-minute timing (seawater coefficient = 0.906–1.040, Holm corrected *P* < 0.018), but not at the 5-minute timing. *Gymnogobius petschiliensis* exhibited no significant variation in this ratio. There were no significant differences in the ratio of floating individuals for any species at either time point.

**Fig. 4.**
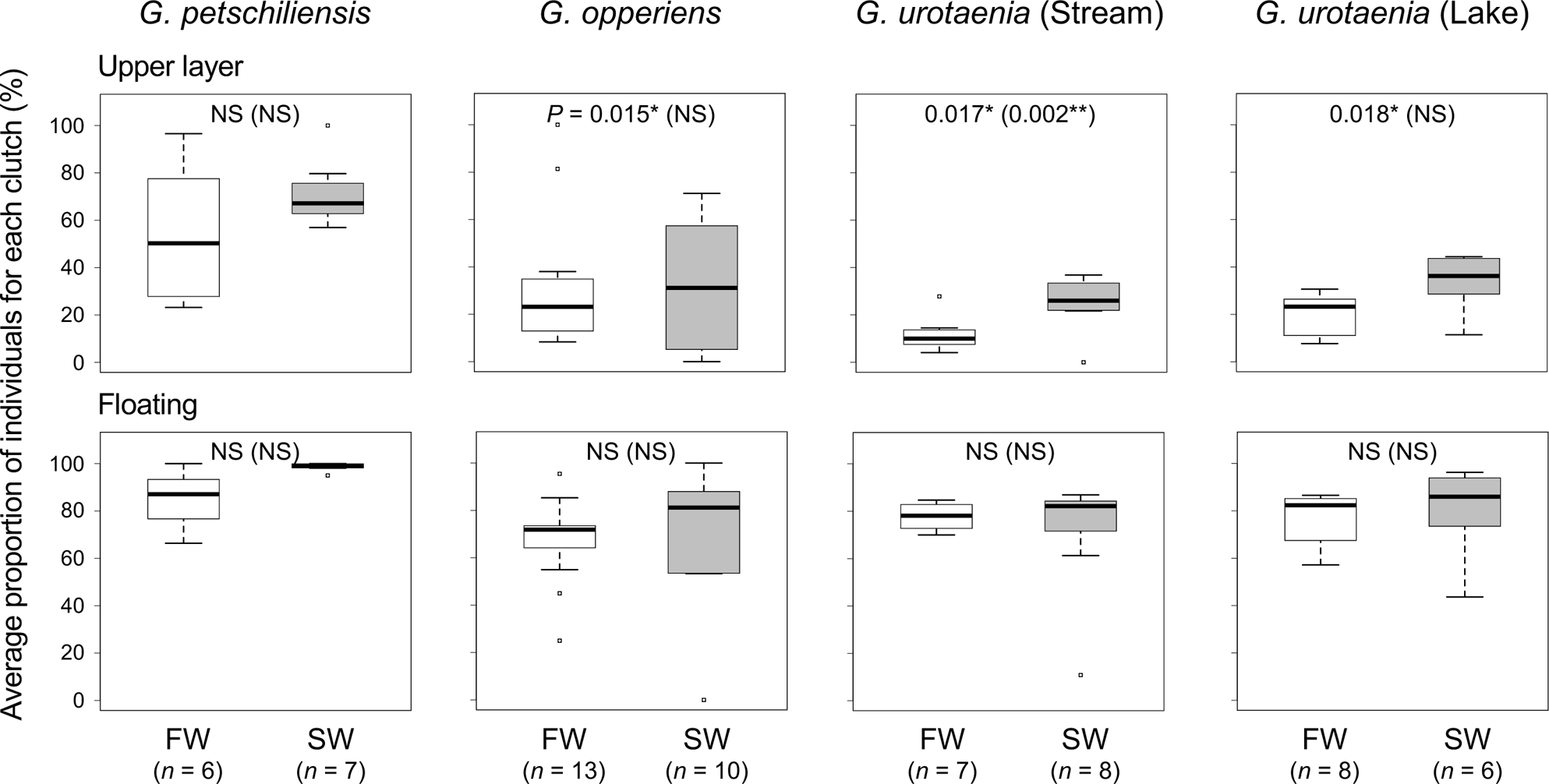
Comparison of swimming layers of *Gymnogobius* larvae between freshwater (FW) and seawater (SW) conditions or among the three targeted species. The sample size below the box plot labels shows the number of clutches from which larval swimming was observed for each salinity group. The *P*-values in the frames indicate the probability that the swimming layers were not different between FW and SW conditions in each species at 5 and 10 minutes after the start of the experiment, with those at the former timing parenthesized (\**P* < 0.05, \*\**P* < 0.01). When the rejection of the null hypothesis was held by the Holm correction process, the *P*-values were replaced by the notation “NS.” The other notations of box plots follow Fig. 2.

The interspecific multiple comparison tests showed that both indices for the swimming layer were significantly higher in *G. petschiliensis* than in the other species at 5- and 10-minute timings (binomial GLMM, coefficient = 0.975–1.858, Holm corrected *P* < 0.030; Table 2), except for the comparison pair with lake *G. urotaenia* at 5-minute timing with a marginal significance (Holm corrected *P* = 0.060). In addition, the ratio of upper swimming individuals was significantly higher in *G. opperiens* than in stream *G. urotaenia* at the 5-minute timing (coefficient = 0.835, Holm corrected *P* = 0.022). No significant differences were found between the other species pairs at any observation time point.

**Table 2.** Results of interspecific comparison of larval swimming behavior in seawater (SW) and freshwater (FW) conditions in the three *Gymnogobius* species, *G. petschiliensis* (*G. pet.*), *G. opperiens* (*G. opp.*), and *G. urotaenia* (*G. uro.*). Populations of *G. urotaenia* were divided into those from rivers (R) and a lake (L)

| Swimming index | Observation timing | Compared species pair | Coefficient (left–right) | Standard error | *z* | *P* | Holm corrected *P* |
| --- | --- | --- | --- | --- | --- | --- | --- |
| Upper layer | 5-minute | *G. pet.*–*G. opp.* | 1.023 | 0.303 | 3.378 | < 0.001*** | 0.003** |
|  |  | *G. pet.*–*G. uro.* (R) | 1.858 | 0.3283 | 5.661 | < 0.001*** | < 0.001*** |
|  |  | *G. pet.*–*G. uro.* (L) | 1.678 | 0.324 | 5.170 | < 0.001*** | < 0.001*** |
|  |  | *G. opp.*–*G. uro.* (R) | 0.835 | 0.311 | 2.686 | 0.007** | 0.022* |
|  |  | *G. opp.*–*G. uro.* (L) | 0.655 | 0.306 | 2.137 | 0.033* | 0.065 |
|  |  | *G. uro.* (R)–*G. uro.* (L) | -0.181 | 0.328 | -0.551 | 0.582 | NS |
|  | 10-minute | *G. pet.*–*G. opp.* | 0.975 | 0.284 | 3.435 | < 0.001*** | 0.002** |
|  |  | *G. pet.*–*G. uro.* (R) | 1.527 | 0.302 | 5.062 | < 0.001*** | < 0.001*** |
|  |  | *G. pet.*–*G. uro.* (L) | 1.423 | 0.301 | 4.730 | < 0.001*** | < 0.001*** |
|  |  | *G. opp.*–*G. uro.* (R) | 0.552 | 0.292 | 1.886 | 0.059 | NS |
|  |  | *G. opp.*–*G. uro.* (L) | 0.448 | 0.291 | 1.537 | 0.124 | NS |
|  |  | *G. uro.* (R)–*G. uro.* (L) | -0.104 | 0.305 | -0.340 | 0.734 | NS |
| Floating | 5-minute | *G. pet.*–*G. opp.* | 1.263 | 0.392 | 3.222 | 0.001** | 0.008** |
|  |  | *G. pet.*–*G. uro.* (R) | 1.156 | 0.405 | 2.853 | 0.004** | 0.022* |
|  |  | *G. pet.*–*G. uro.* (L) | 0.989 | 0.406 | 2.433 | 0.015* | 0.060 |
|  |  | *G. opp.*–*G. uro.* (R) | -0.107 | 0.316 | -0.339 | 0.735 | NS |
|  |  | *G. opp.*–*G. uro.* (L) | -0.274 | 0.325 | -0.844 | 0.399 | NS |
|  |  | *G. uro.* (R)–*G. uro.* (L) | -0.167 | 0.343 | -0.487 | 0.626 | NS |
|  | 10-minute | *G. pet.*–*G. opp.* | 1.556 | 0.414 | 3.760 | < 0.001*** | 0.001** |
|  |  | *G. pet.*–*G. uro.* (R) | 1.224 | 0.430 | 2.849 | 0.004** | 0.022* |
|  |  | *G. pet.*–*G. uro.* (L) | 1.152 | 0.431 | 2.673 | 0.008** | 0.030* |
|  |  | *G. opp.*–*G. uro.* (R) | -0.332 | 0.325 | -1.022 | 0.307 | NS |
|  |  | *G. opp.*–*G. uro.* (L) | -0.404 | 0.331 | -1.221 | 0.222 | NS |
|  |  | *G. uro.* (R)–*G. uro.* (L) | -0.072 | 0.351 | -0.204 | 0.838 | NS |

**Note 1:** The formula of GLMM was as follows: Swimming index of each individual (0/1; assumed to follow a binomial distribution) ~ Salinity group + Species. Salinity group was a categorical variable (i.e., FW or SW). Population and parent IDs of each larva were also considered as random effects.
**Note 2:** \**P* < 0.05, \*\**P* < 0.01, \*\*\**P* < 0.001.
**Note 3:** If the null hypothesis could not be rejected in the Holm method, the *P*-values of the subsequent tests were not corrected (indicated with "NS").

## 4. Discussion

This study tested whether the ability of larvae to regulate body specific gravity (SG) is linked to the occurrence of freshwater resident (FWR) forms in amphidromous *Gymnogobius* species. Contrary to our expectations, the obtained results consistently suggest that all targeted species have a high ability to modulate body SG, regardless of the presence or absence of FWR populations in species. This section first discusses the SG modulation flexibility of *Gymnogobius* larvae by comparing it with those of other fish species and the physiological traits that enhance this ability in a physiological context. Second, we discuss when the SG flexibility was acquired and how it has contributed to freshwater colonization in an evolutionary context.

### 4.1 SG regulation performance

Generally, the loss of larval neutral buoyancy results in high energy consumption to maintain a vertical position (Czesny et al., 2005; Woolley and Qin, 2010), low anti-predatory performance (Czesny et al., 2005), and morphological abnormalities (Chatain, 1989; Schwebel et al., 2018). For the larvae of amphidromous *Gymnogobius* species, which prey on planktonic copepods and amphipods drifting in the middle and surface layers (Dotu, 1955), the loss of neutral buoyancy should negatively affect their feeding performance and growth. Their flexible modulation of body SG would strongly facilitate normal growth and survival, even when the environmental SG is low.

To our knowledge, *Gymnogobius* larvae showed higher SG modulation ability (approximately 1.001–1.030; Fig. 2) than any other aquatic organisms that have ever been studied. For example, the SG of newly hatched larvae of sicydiine gobies ranges from 1.013 to 1.036, which is usually higher than that of ambient brackish water or seawater (Iida et al., 2010, 2017). Furthermore, the ranges of SGs at any specific life history stage are narrower than the above values because of irreversible changes due to growth in sicydiine gobies. Similarly, body SG ranges tend to be narrower than the SG ranges of ambient waters in several other goby species (Iida et al., 2017), Baltic cod (Nissling and Vallin, 1996), and black sea bream (Kitajima et al., 1994). Few studies have investigated the SG modulation ability in aquatic invertebrate larvae. However, as they do not have a swimbladder, the modulation seems difficult for them; their body SGs appear to be highly species-specific (<0.005 in the SG range per species) (Moore et al., 1997; Davenport and Healy, 2006). Thus, to the best of our knowledge, the finding for *Gymnogobius* species is the first report that fish (or any aquatic organism) can maintain their neutral buoyancy in both freshwater and seawater environments by flexibly adjusting their body SG, even during the early larval stage.

A swimbladder is suggested to contribute considerably to the flexible SG regulation in *Gymnogobius* larvae because their swimbladder size was much larger in freshwater than in seawater. Generally, the swimbladder of early larval fish has poorly function; for example, it starts to function after initial air intake at the age of 5–10 days in red sea bream (Kitajima et al., 1993), 6–11 days in Australian bass (Battaglene and Talbot, 1990), and 5–11 days in mandarin fish (He et al., 2022). Also, in many other teleost species, larvae do not uptake air into their swimbladder until approximately one week (at least three days) after hatching (striped bass: Martin-Robichaud and Peterson, 1998; striped trumpeter: Trotter et al., 2001; Pacific bluefin tuna: Takashi et al., 2006; blacktip grouper: Kawabe and Kimura, 2008; zebrafish: Winata et al., 2009; burbot: Palińska-Żarska et al., 2014). In contrast, all three *Gymnogobius* species had already inflated their swimbladders in most 3-day-old larvae (Fig. 3A, B). Furthermore, the inflation often occurred immediately after hatching (Fig. 3B), indicating that the swimbladder begins to function earlier in the *Gymnogobius* gobies than in other previously reported fish species.

Although this study did not find considerable differences in the swimming layers between the freshwater and seawater conditions, the data might not effectively reflect those in the wild due to the small experimental tank. Further experiments using larger aquariums and field surveys are needed to confirm the swimming layer in natural conditions.

### 4.2 Pre-adaptation to low SG environment

The highly functional swimbladders of early larvae in the *Gymnogobius* species may have been pre-adaptively acquired prior to larval freshwater colonization. This study showed that the obligate amphidromous *G. petschiliensis* and *G. opperiens* can inflate their swimbladders at the early larval stage, similar to the freshwater-dependent amphidromous *G. urotaenia*, which has derived FWR types. Phylogenetic studies suggest that *Gymnogobius* originated from a marine or estuarine ancestral species, and that *G. petschiliensis* and *G. opperiens* were derived earlier than *G. urotaenia* (Aizawa et al., 1994; Harada et al., 2002; Ito et al., 2022). This suggests that the obligate amphidromous species were directly derived from marine or estuarine species, and thus the high SG modulation ability was not inherited from any strictly freshwater species. The ancestor of the *Gymnogobius* species is presumed to have inhabited shallow coastal or estuarine areas, where salinity frequently fluctuated due to stream flow, rain, and intense temperature fluctuations (Harada, 2005; Ito et al., 2022). Such environmental conditions experienced by ancestral species are inferred to have evolutionarily enhanced the flexibility of body SG modulation, allowing goby larvae to effectively maintain neutral buoyancy, even when unexpectedly trapped in freshwater habitats.

The evolutionary process of the SG modulation ability may differ from that of osmoregulatory flexibility, another important physiological trait (McCairns and Bernatchez, 2010; Schultz and McCormick, 2013). The larval osmoregulatory capacity under freshwater conditions appears to be higher in the freshwater-dependent amphidromous *G. urotaenia* than in *G. petschiliensis* and *G. opperiens* (Oto et al., 2022). This interspecific difference may be related to the difficulty in evolving and maintaining a highly plastic osmoregulatory mechanism because of the underdeveloped osmoregulatory organs in early larvae (Kaneko et al., 2002; Varsamos et al., 2005) and the trade-offs between osmoregulatory ability in hypertonic and hypotonic environments (Velotta et al., 2015). Thus, this study shows a case in which different freshwater-adapted traits may have evolved asynchronously during freshwater colonization in a fish group. However, since this study used a limited number of species in a clade, the expansion of taxon sampling is needed to verify the scenario of pre-adaptation to low SG environments.

### 4.3 Importance of flexible SG regulation in life-history divergence

The laboratory observations revealed that the larval swimming layer was not greatly affected by salinity in either species. On the other hand, the swimming layers differed among species; in particular, *G. petschiliensis* larvae strongly preferred water surfaces, regardless of salinity. These species-specific larval swimming layers may reflect variations in dispersal strategies.

Alteration of the vertical larval position in the water column can change their arrival points on a scale of tens to hundreds of kilometers (Cowen et al., 2000; Bradbury and Snelgrove, 2001; Secor, 2015); that is, larvae near the surface can be transported over longer distances than those near the bottom by catching the outflow from rivers (Dame and Allen, 1996). Therefore, the preference for a swimming layer near the water surface in *G. petschiliensis* larvae might result in greater extent of dispersal in this species. The link between dispersal and larval characteristics can be further examined by analyzing the otolith microchemical composition and spatial genetic structure.

## 5. Conclusion

In physiological and evolutionary studies on freshwater colonization of fish, osmoregulatory and/or nutritional problems have been examined as factors limiting larval survival in freshwater habitats (Ishikawa et al., 2019; Oto et al., 2022), but the maintenance of neutral buoyancy under low ambient specific gravity (SG) has received little attention. This study tested the hypothesis regarding the relationship between the occurrence of freshwater resident populations and larval body SG modulation ability in an amphidromous goby group through interspecific comparisons. The results rejected the initial hypothesis but consistently suggested that SG modulation ability is a pre-adaptive rather than a newly evolved trait that has enabled residence in freshwater habitats. This evolutionary process of SG modulation ability seems to differ from that of osmoregulatory flexibility, which correlates with the occurrence of freshwater resident populations in the target goby group (Oto et al., 2022). The generality of the pre-adaptation scenario to low environmental SG should be tested by examining the relationships between the larval SG modulation ability and the occurrence of freshwater resident populations across wider taxonomic groups in a phylogenetic context.

## Acknowledgements

We thank S. Kunimatsu for his help with the molecular species identification and members of Laboratory of Animal Ecology of Kyoto University for their helpful comments on the study design.

## Ethical approval

All applicable international, national, and/or institutional guidelines for the care and use of animals were followed.

## Data availability

The registration of the raw data for the analyses of body specific gravity, swimbladder size, and swimming layer is in progress in DRYAD (https://datadryad.org).

## Author contributions

Y.O. and K.W conceived the presented idea. Y.O. performed the fish sampling and experiments. Y.O. and K.W wrote the manuscript.

