## Supplementary Figs. S1-S3 for "Does larval ability to modulate body buoyancy explain successful colonization of freshwater environments by diadromous gobies?"

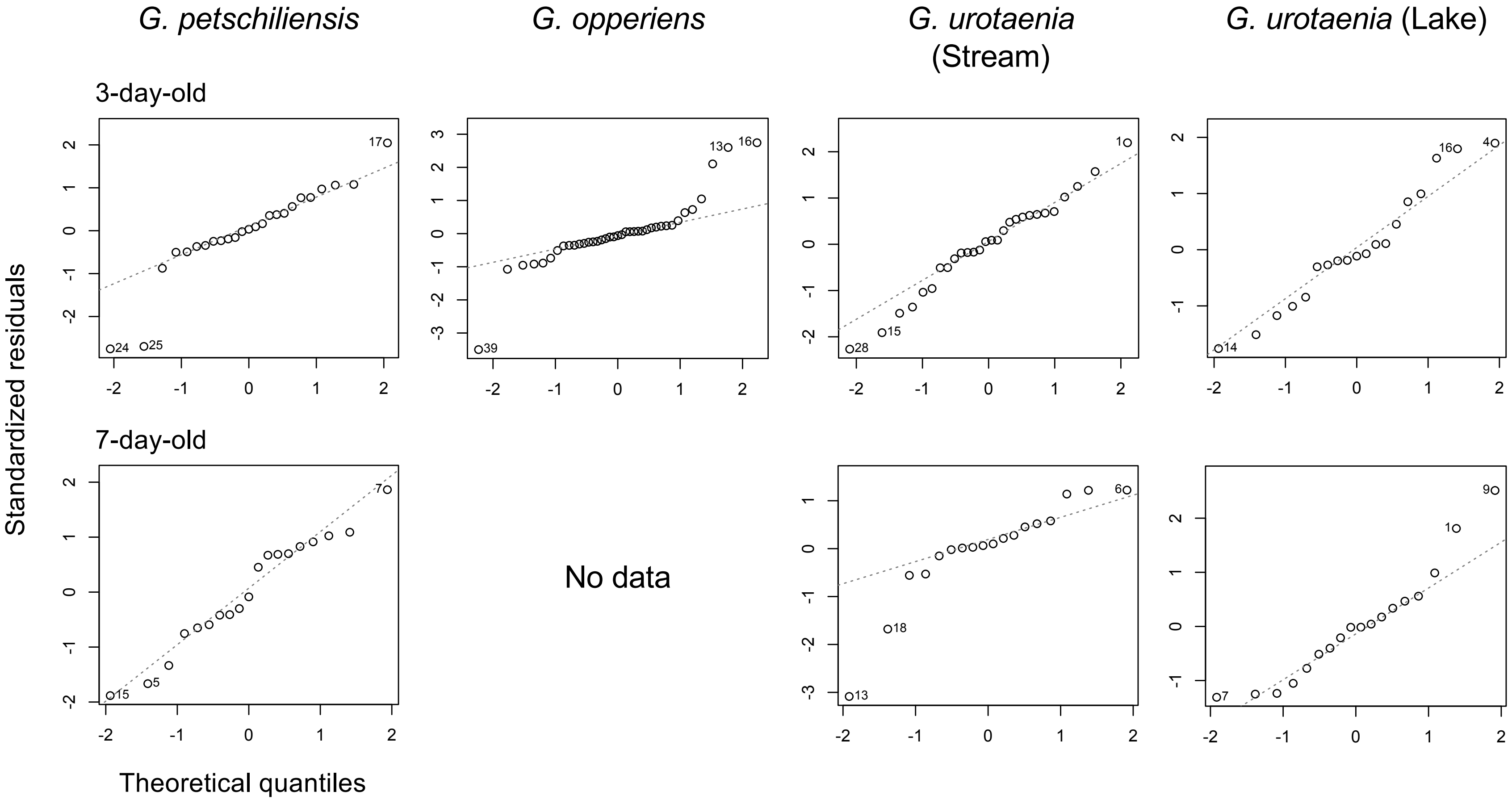


**Fig. S1** Quantile-quantile plots to examine the normality of the response variable of swimbladder size in multiple regression analysis. Ideally, all plots are aligned on the dotted line when the variable follows Gaussian distribution. Note that the random effects included in the linear mixed model for the main analysis (population and parent IDs) are not considered in this normality examination.

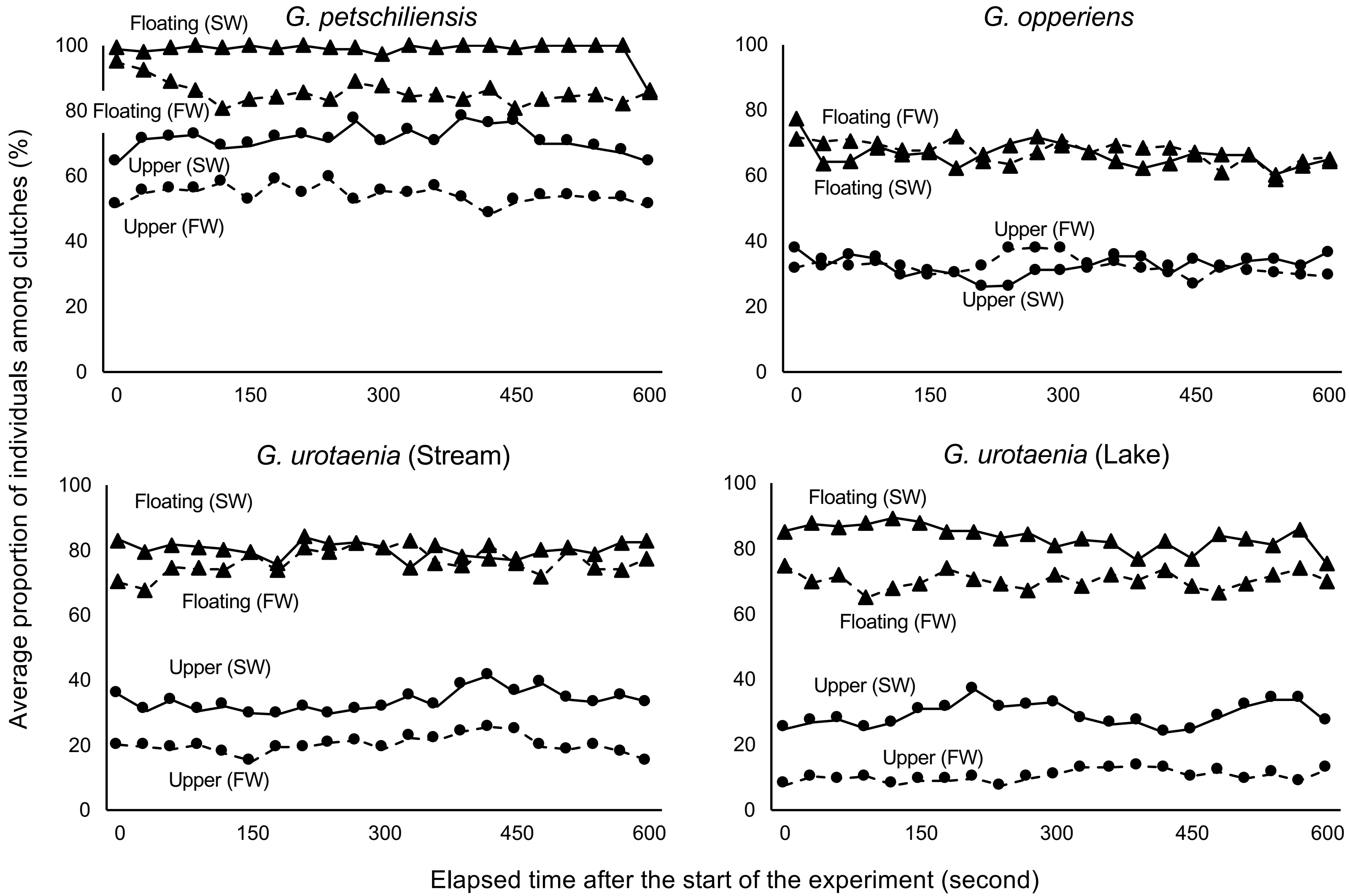


**Fig. S2** The transition of swimming layer during a 10-minute experiment (clutches pooled). The solid and dashed lines indicate seawater (SW) and freshwater (FW) treatments, respectively. The circle and triangle plots indicate the ratios of individuals swimming in the upper-layer section and floating (not sinking to the bottom) in the experimental tank, respectively. The values at each time point represent the means for each salinity group of each species. The number of egg clutches (rearing bottles) of each species was as follows: *G. petschiliensis* SW, $n = 7$, FW, 6; *G. opperiens* SW, 10, FW, 13; *G. urotaenia* (Stream) SW, 7, FW, 8; *G. urotaenia* (Lake) SW, 6, FW, 8. There were no significant time-dependent variations in both indices of the swimming layer in all species (binomial GLMM for each species, Holm-corrected $P > 0.229$). The formula of this GLMM analysis was as follows; binomial variable of swimming layer ~ glmer (Salinity + Elapsed time since the start of the experiment + (1|Parent) + (1|Population)). The Holm correction was conducted per salinity group to consider the effect of the repetition of intraspecific comparisons.

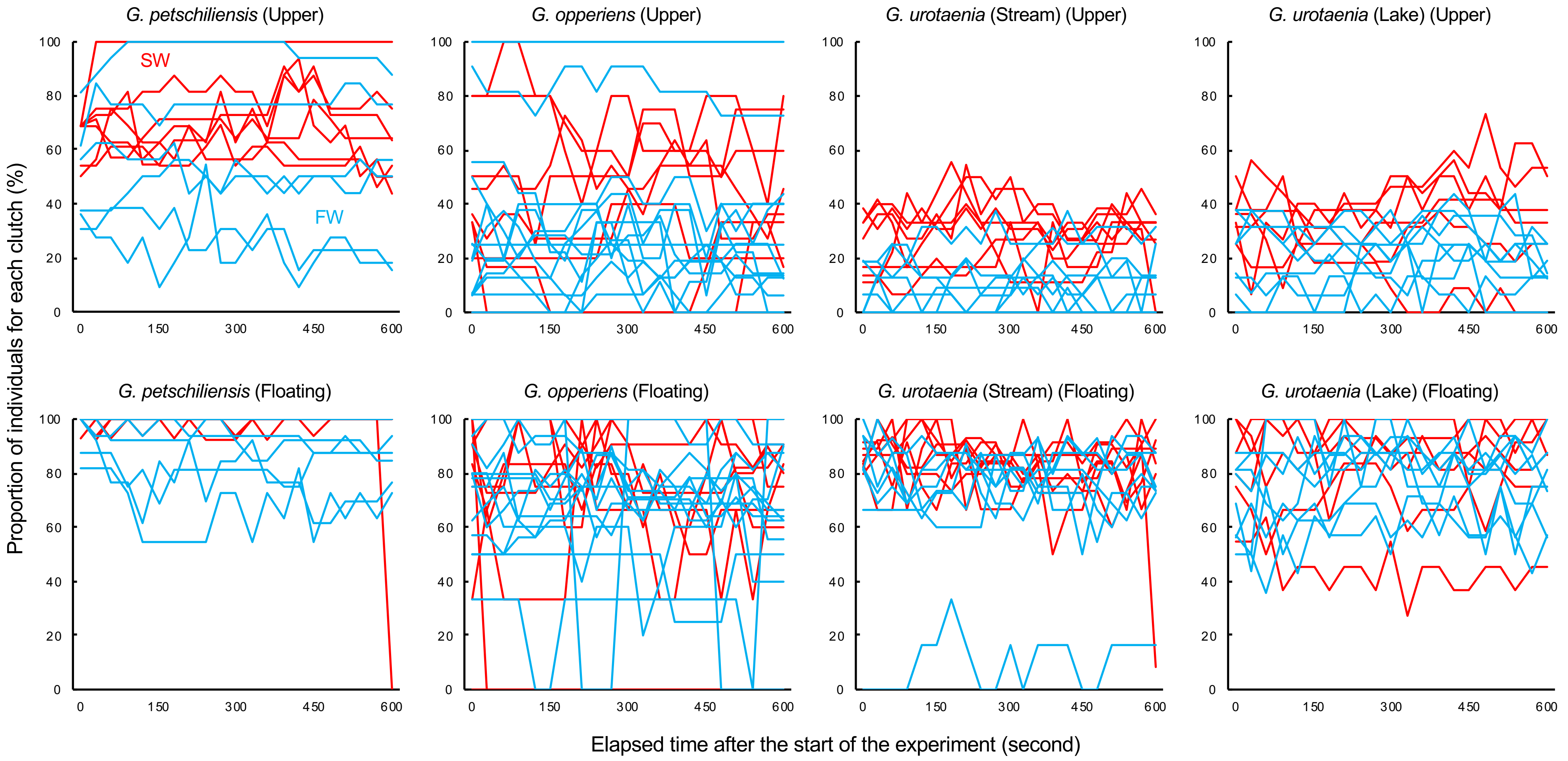


**Fig. S3** The transition of swimming layer during a 10-minute experiment (clutches not pooled). The red and blue lines indicate seawater (SW) and freshwater (FW) treatments, respectively. The upper and lower graphs show the ratios of individuals swimming at the upper-layer section and floating (i.e., not sunk to the bottom) in the experimental tank, respectively. For the number of egg clutches (rearing bottles) of each species, please see Fig. S2.
